# BceAB-type antibiotic resistance transporters appear to act by target protection of cell wall synthesis

**DOI:** 10.1101/835702

**Authors:** Carolin M Kobras, Hannah Piepenbreier, Jennifer Emenegger, Andre Sim, Georg Fritz, Susanne Gebhard

## Abstract

Resistance against cell wall-active antimicrobial peptides in bacteria is often mediated by transporters. In low GC-content Gram-positive bacteria, a wide-spread type of such transporters are the BceAB-like systems, which frequently provide a high level of resistance against peptide antibiotics that target intermediates of the lipid II cycle of cell wall synthesis. How a transporter can offer protection from drugs that are active on the cell surface, however, has presented researchers with a conundrum. Multiple theories have been discussed, ranging from removal of the peptides from the membrane, internalisation of the drug for degradation, to removal of the cellular target rather than the drug itself. To resolve this much-debated question, we here investigated the mode of action of the transporter BceAB of *Bacillus subtilis*. We show that it does not inactivate or import its substrate antibiotic bacitracin. Moreover, we present evidence that the critical factor driving transport activity is not the drug itself, but instead the concentration of drug-target complexes in the cell. Our results, together with previously reported findings, lead us to propose that BceAB-type transporters act by transiently freeing lipid II cycle intermediates from the inhibitory grip of antimicrobial peptides, and thus provide resistance through target protection of cell wall synthesis. Target protection has so far only been reported for resistance against antibiotics with intracellular targets, such as the ribosome. However, this mechanism offers a plausible explanation for the use of transporters as resistance determinants against cell wall-active antibiotics in Gram-positive bacteria where cell wall synthesis lacks the additional protection of an outer membrane.

## INTRODUCTION

The bacterial cell wall and its biosynthetic pathway, the lipid II cycle, are important targets for antibiotics, especially in Gram-positive bacteria that lack the protective layer of the outer membrane. Cell wall-targeting drugs include antimicrobial peptides (AMPs), which bind to cycle intermediates and prevent biosynthetic enzymes from carrying out the next reaction (1). It is hardly surprising that bacteria have developed a plethora of strategies to protect themselves against such antibiotic attack. Among the many known resistance mechanisms, a common strategy is the production of ATP-binding cassette (ABC) transporters that presumably remove AMPs from their site of action (2, 3). A major group of these are the BceAB-type transporters, which are found in many environmental and pathogenic species of the phylum Firmicutes (4). The eponymous and to date best-characterised system is BceAB of *Bacillus subtilis* (5). BceAB-type transporters comprise one permease (BceB) and two ATPases (BceA) (BceA, 6). The permeases consist of ten transmembrane helices and a large extracellular domain that is thought to contain the ligand binding region of the transporter (7, 8). Transporter production is regulated via a two-component regulatory system (TCS) consisting of a histidine kinase (BceS) and a response regulator (BceR) (BceR, 5, 7). A striking feature of these systems is that signalling is triggered by the activity of the transporter itself (9). Due to this flux-sensing strategy, signalling is directly proportional to transport activity, and the transporter effectively autoregulates its own production (Fig. 1A).

**Fig. 1:**
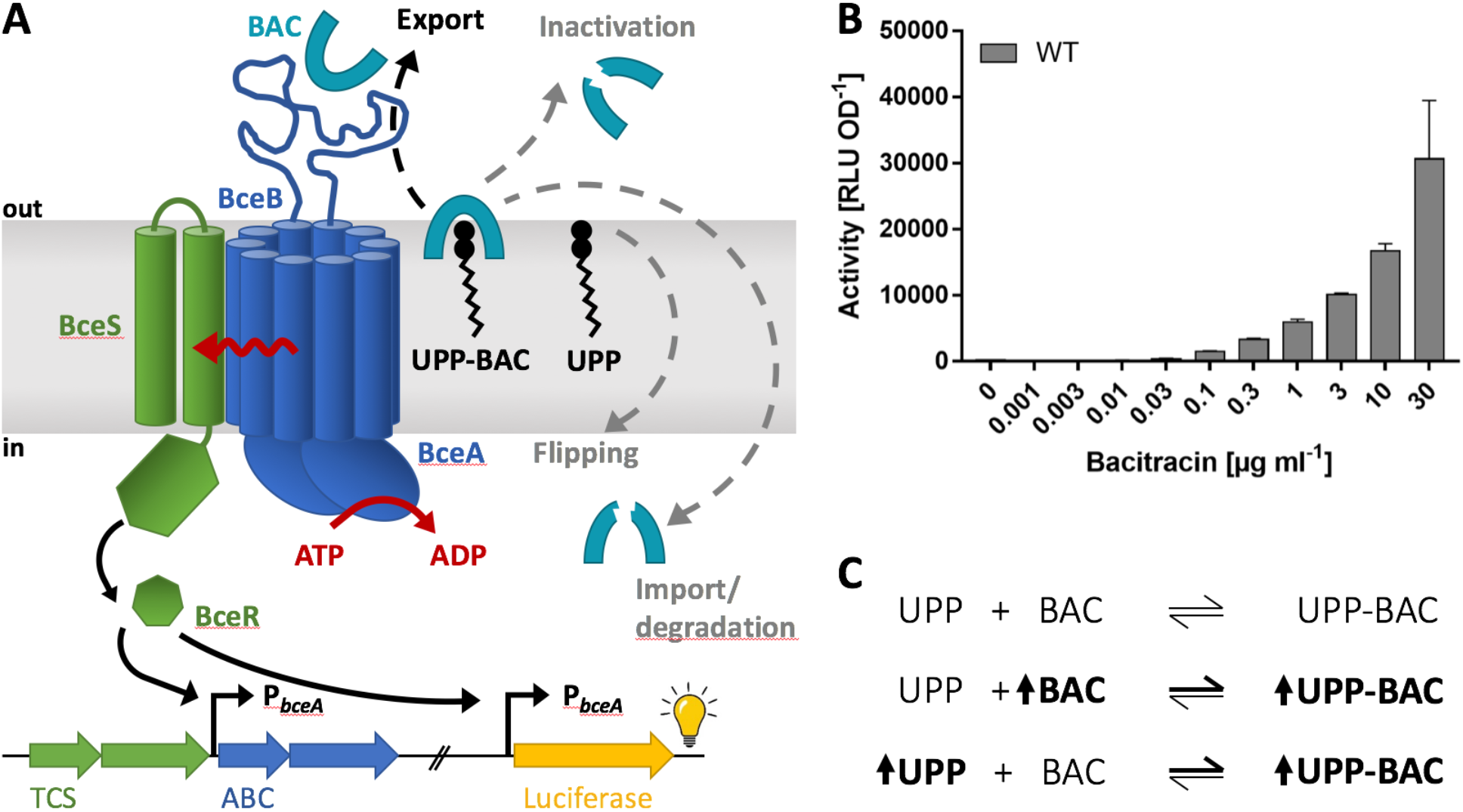
Antibiotic resistance and flux-sensing by BceAB. **A:** Schematic of the BceAB-BceRS resistance system. The transporter BceAB confers resistance against bacitracin (BAC), which acts by binding its cellular target UPP. The different debated mechanisms for resistance by BceAB are indicated by dashed arrows (see text for details). Flux-sensing communicates the transport activity of BceAB to the kinase BceS (red wave arrow), causing activation of BceR, which induces transcription from the target promotor P_*bceA*_. This results in increased production of BceAB, and therefore adjusted levels of resistance. As signalling is directly proportional to BceAB activity, we can use the target promotor P_*bceA*_ fused to a luciferase reporter to monitor transport activity. TCS, genes encoding the two-component regulatory system BceRS; ABC, genes encoding the resistance transporter BceAB. **B:** Using luciferase activity as a proxy, BceAB activity of wild-type *B. subtilis* W168 carrying the P_*bceA*_-*lux* reporter fusion (WT, SGB73) was determined following 25-35 min challenge of exponentially growing cells with sub-inhibitory concentrations of bacitracin. All data are depicted as mean ± standard deviation of at least three biological replicates. **C:** Binding reaction between free bacitracin and its cellular target UPP. The change in concentration of UPP-bacitracin complexes (UPP-BAC) through manipulation of either bacitracin or UPP concentrations is indicated by bold font and upward-facing arrows.

BceAB confers resistance against the AMPs bacitracin, mersacidin, actagardine and plectasin, of which bacitracin binds the lipid II cycle intermediate undecaprenyl pyrophosphate (UPP), while the others bind lipid II itself (5, 8). Considering the location of the AMPs’ targets on the extracellular side of the cytoplasmic membrane, it is not immediately obvious how a membrane-embedded transporter can provide effective protection from these drugs. The mode of action of BceAB-type transporters has therefore been the subject of much debate (Fig. 1A). When first described, the *B. subtilis* system was named Bce for <u>b</u>a<u>c</u>itracin <u>e</u>fflux (5), although no evidence for the direction of transport was available. The assumption of export was based on the suggested self-protection mechanism of the unrelated transporter BcrAB in the bacitracin producer *B. licheniformis* ATCC10716 (10, 11). BcrAB was thought to work as a ‘hydrophobic vacuum cleaner’ to remove the antibiotic from the membrane, akin to the human multidrug resistance transporter P-glycoprotein (12, 13). Later, BceAB was speculated to instead import bacitracin into the cytoplasm for subsequent degradation, again without direct experimental evidence (7). More recently, the transporter was proposed to act as a UPP flippase (14). In this scenario, BceAB would confer resistance by transporting UPP across the membrane to the cytoplasmic face, thereby removing the cellular target for bacitracin rather than transporting bacitracin itself. In the presence of bacitracin, BceAB was hypothesised to be inhibited by UPP-bacitracin complexes (UPP-BAC), which in turn should activate signalling through the BceRS two-component system to adjust BceAB levels in the cell (14). This model offered a neat explanation of the available data on bacitracin resistance, but could not explain how the same transporter can confer resistance against AMPs that target lipid II instead of UPP.

Since then, we have shown that BceB is able to bind bacitracin *in vitro* (6). Without excluding the possibility of BceAB interacting with the UPP-BAC complex, this finding suggested that BceAB-like transporters directly interacted with the AMP and that the AMP is at least part of the physiological substrate. Moreover, the computational model used to establish the flux-sensing mechanism for signalling within the Bce system was based on recognition of UPP-BAC complexes by the transporter and removal of bacitracin from the complex (9). Although the model did not specify a particular direction of transport, such a mechanism was most in line with the initial hydrophobic vacuum cleaner hypothesis (5). Resistance in this scenario is conferred by BceAB recognising target-AMP complexes in the membrane, removing the antibiotic and releasing it into the extracellular milieu. This frees the target from the inhibitory action of the antibiotic and allows the next step of cell wall synthesis to proceed.

Considering the relevance of BceAB-like systems among Firmicutes bacteria, we here set out to address the controversial question on their mode of action and how a transporter can provide effective protection against cell surface-active antibiotics. Using a peptide release assay, we exclude that BceAB acts by import or inactivation of bacitracin. Based on the discovery that signalling within the Bce system is directly proportional to transport activity, we established a promoter-reporter assay as a proxy for transport activity. Our results show that the critical variable in determining transport activity of BceAB is bacitracin in complex with its cellular target UPP, rather than bacitracin or the lipid carrier alone. Taking together the findings of this study and the literature, we conclude that BceAB-type transporters appear to transiently free their cellular target from the inhibitory grip of the AMP and provide resistance via target protection of cell wall synthesis.

## METHODS

### Bacterial strains and growth conditions

All strains used in this study are given in Table 1. *E. coli* and *B. subtilis* strains were routinely grown at 37 °C with agitation (180 rpm) in lysogeny broth (LB) medium. Solid media contained 1.5% (w/v) agar. Selective media contained ampicillin (100 μg ml^−1^), chloramphenicol (5 μg ml^−1^), kanamycin (10 μg ml^−1^), spectinomycin (100 μg ml^−1^), tetracycline (10 μg ml^−1^) or erythromycin (1 μg ml^−1^) with lincomycin (25 μg ml^−1^; macrolide-lincosamide-streptogramin B; mls). For full induction of the promoter P_*xylA*_, xylose was added to a final concentration of 0.2 % (w/v). Bacterial growth was routinely monitored as optical density at 600 nm wavelength (OD_600_) measured spectrophotometrically in cuvettes of 1 cm light path length.

**Table 1:** Plasmids and bacterial strains used in this study.

| Name | Genotype and description<sup>a</sup> | Source |
| --- | --- | --- |
| **Plasmids** |  |  |
| pAH328 | Vector for transcriptional promoter fusions to *luxABCDE*; integrates in *B. subtilis* *sacA*; Amp<sup>r</sup>, Cm<sup>r</sup> | (52) |
| pBS2E | Empty vector; integrates in *B. subtilis* *lacA*; Amp<sup>r</sup>, mls<sup>r</sup> | (16) |
| pBS3Elux | Vector for transcriptional promoter fusions to *luxABCDE*; integrates in *B. subtilis* *sacA*; Amp<sup>r</sup>, Mls<sup>r</sup> | (18) |
| pSB1A3 | Empty BioBrick standard cloning vector for *E. coli*; Amp<sup>r</sup> | Registry of Standard Biological Parts |
| pSDLux101 | pAH328 harbouring a transcriptional P<sub>bceA</sub>-*luxABCDE* fusion | (17) |
| pSDLux102 | pAH328 harbouring a transcriptional P<sub>psdA</sub>-*luxABCDE* fusion | This study |
| pJNESB101 | pSB1A3 harbouring *B. subtilis* *bcrC* in BioBrick format | This study |
| pJNE2E01 | pBS2E harbouring a transcriptional P<sub>xyIA</sub>-*bcrC* fusion assembled according to the BioBrick RFC25 cloning standard | This study |
| pNT2E01 | pBS2E harbouring P<sub>xyIA</sub>-*bceAB* assembled according to the BioBrick RFC10 cloning standard | (9) |
| pMG3Elux1 | pBS3Elux harbouring a transcriptional P<sub>bceA</sub>-*luxABCDE* fusion | This study |
| ***B. subtilis* strains** |  |  |
| W168 | Wild type, *trpC2* | Laboratory stock |
| TMB035 ($\Delta bceAB$) | W168 *bceAB*::kan; Kan<sup>r</sup> | (7) |
| TMB297 ($\Delta bcrC$) | W168 *bcrC*::tet; Tet<sup>r</sup> | (7) |
| TMB713 ($\Delta bceAB \Delta bcrC$) | W168 *bceAB*::kan *bcrC*::tet; Kan<sup>r</sup>, Tet<sup>r</sup> | (24) |
| HB13350 | W168 *ytpB*::spec; Spec<sup>r</sup> | (14) |
| HB13438 | W168 *menA*::kan *amyE*::P<sub>spac(hy)</sub>-*menA*; Kan<sup>r</sup>, Cm<sup>r</sup> | (14) |
| SGB73 | W168 *sacA*::pSDLux101; Cm<sup>r</sup> | (9) |
| SGB74 | W168 *sacA*::pSDLux102; Cm<sup>r</sup> | This study |
| SGB218 | W168 *bceAB*::kan *sacA*::pSDLux101 *lacA*::pNT2E01; Kan<sup>r</sup>, Cm<sup>r</sup>, Mls<sup>r</sup> | (9) |
| SGB243 | W168 *lacA*::pJNE2E01; Mls<sup>r</sup> | This study |
| SGB649 | W168 *bcrC*::tet *sacA*::pSDLux101; Tet<sup>r</sup>, Cm<sup>r</sup> | This study |
| SGB677 | W168 *bceAB*::kan *bcrC*::tet *sacA*::pSDLux101 *lacA*::pNT2E01; Kan<sup>r</sup>, Tet<sup>r</sup>, Cm<sup>r</sup>, Mls<sup>r</sup> | This study |
| SGB681 | W168 *bcrC*::tet *sacA*::pSDLux102; Tet<sup>r</sup>, Cm<sup>r</sup> | This study |
| SGB758 | W168 *sacA*::pSDLux101 *lacA*::pJNE2E01; Cm<sup>r</sup>, Mls<sup>r</sup> | This study |
| SGB873 | W168 *menA*::kan *amyE*::P<sub>spac(hy)</sub>-*menA* *ytpB*::spec; Kan<sup>r</sup>, Cm<sup>r</sup>, Spec<sup>r</sup> | This study |
| SGB927 | W168 *sacA*::pMG3Elux1; Mls<sup>r</sup> | This study |
| SGB929 | W168 *menA*::kan *amyE*::P<sub>spac(hy)</sub>-*menA* *ytpB*::spec *sacA*::pMG3Elux1; Kan<sup>r</sup>, Cm<sup>r</sup>, Spec<sup>r</sup>, Mls<sup>r</sup> | This study |
| SGB974 | W168 *sacA*::pSDlux102 *lacA*::pJNE2E01; Cm<sup>r</sup>, Mls<sup>r</sup> | This study |

<sup>a</sup> Amp<sup>r</sup>, ampicillin resistance; Cm<sup>r</sup>, chloramphenicol resistance; Kan<sup>r</sup>, kanamycin resistance; Mls<sup>r</sup>, macrolide, lincosamide and streptogramin B resistance; Tet<sup>r</sup>, tetracycline resistance, Spec<sup>r</sup>, spectinomycin resistance

### Strain construction and molecular cloning

All plasmids used in this study are listed in Table 1; primer sequences are given in Table 2. *B. subtilis* transformations were performed using a modified version of the Paris protocol (15). Overnight cultures of recipient strains were grown in 500 μl Paris medium (6.1 mM K_2_HPO_4_, 4.4 mM KH_2_PO_4_, 0.4 mM trisodium citrate, 1 % (w/v) glucose, 20 mM potassium L-glutamate, 0.1 % (w/v) casamino acids, 3 mM MgSO_4_, 25 μg ml^−1^ tryptophan, 8 μM ferric ammonium citrate) at 37 °C with aeration (180 rpm). Day cultures (500 μl) were inoculated 1:50 in fresh, pre-warmed Paris medium and grown for three hours (37 °C, 180 rpm). To each culture, 50 μl of isolated genomic DNA (gDNA) of the donor strain, or 0.5-1 μg of isolated plasmid DNA were added. Transformation cultures were grown for five more hours and plated on selective media. For mls or chloramphenicol resistance, cultures were pre-induced for one hour at 1:40 of the final concentration of the respective antibiotic. Donor strain gDNA was isolated by mixing an overnight culture of the donor 1:1 with SC buffer (150 mM NaCl, 10 mM sodium citrate, pH 7.0) and harvesting the cells by centrifugation (5 min, 1300 × *g*). The pellet was resuspended in SC buffer and incubated with lysozyme at 37 °C for 15 minutes. The solution was mixed 1:1 with 5 M NaCl and passed through a 0.45 μm syringe-driven filter. Plasmid DNA was isolated from *E. coli* using conventional mini-prep kits.

**Table 2:**
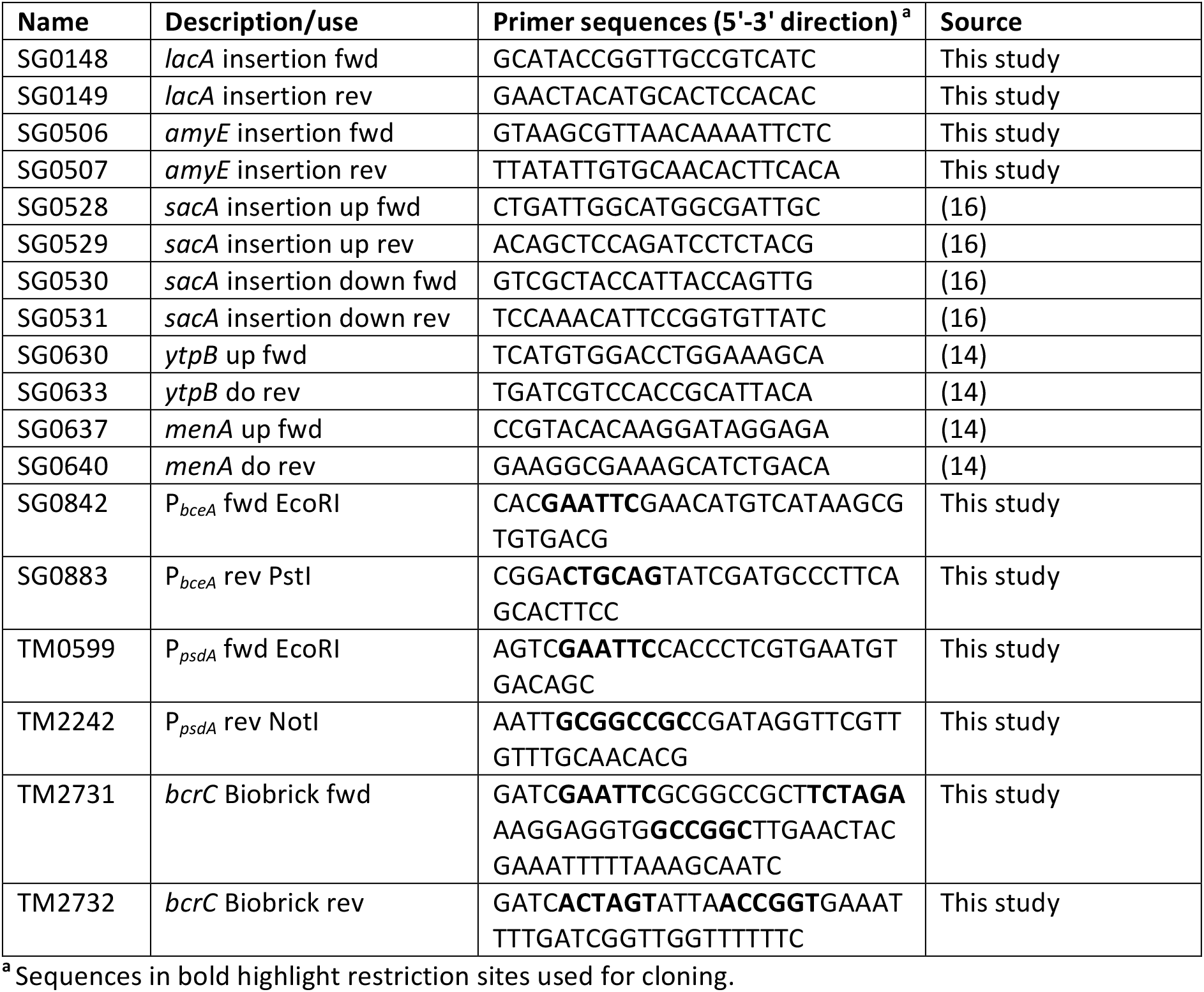
Primers used in this study.

To create a construct for inducible expression of *bcrC*, the gene was PCR-amplified from *B. subtilis* W168 using primers TM2731 and TM2732, which incorporated prefix and suffix, respectively, of the modified ‘Freiburg standard’ of BioBrick cloning described previously (16). The resulting fragment was cloned into pSB1A3 via the EcoRI and PstI restriction sites (pJNESB101). The *bcrC* gene was then re-excised using XbaI and PstI. Assembly with an EcoRI/SpeI fragment of the BioBrick carrying the xylose-inducible promoter P_*xylA*_ (16) into EcoRI/PstI digested pBS2E resulted in the inducible P_*xylA*_-*bcrC* construct pJNE2E01. A transcriptional P_*psdA*_-*lux* reporter construct (pSDlux102) was created by PCR amplification of the promoter region of *psdAB* of *B. subtilis* using primers TM0599 and TM2242, and ligation with pAH328 via EcoRI and NotI restriction sites. The existing P_*bceA*_-*luxABCDE* reporter (pSDlux101; (17) was re-constructed in vector pBS3Elux (18), which contains an mls resistance marker instead of chloramphenicol. This was achieved by PCR amplification of the promoter fragment with primers SG843 and SG883, and cloning via EcoRI and PstI sites, resulting in plasmid pMG3Elux1.

### Determination of the minimal inhibitory concentration

The susceptibility of *B. subtilis* strains to bacitracin was determined using the minimal inhibitory concentration (MIC) determined by broth micro-dilutions. For this, two-fold serial dilutions of Zn^2+^-bacitracin were prepared in 2 ml of Mueller-Hinton medium and inoculated 1:500 from overnight cultures grown in the same medium. For higher resolution, in some instances defined concentrations of Zn^2+^-bacitracin were added directly to each culture. Cultures were incubated overnight (37 °C, 180 rpm) and examined for growth after 24 h. The MIC was determined as the lowest concentration at which no visible growth was detected. All experiments were performed in at least biological triplicates, and mean values and standard deviations were calculated to report the data.

### Bacitracin uptake assays

Bacitracin uptake was assayed with slight modification to previously described protocols (19, 20). Overnight cultures were diluted 1:500 in 100 ml LB supplemented with 1 % (w/v) fructose. To induce BceAB production in the wild type, 1 μg ml^−1^ bacitracin was added at the time of inoculation. The cultures were incubated for 3.5-4.75 h at 37 °C (200 rpm) until they reached an OD_600_ of 1.0-2.0. Cells were harvested by centrifugation (4000 × *g*, 10 minutes, room temperature) and washed twice with 50 mM potassium phosphate (pH 7-7.5) and 100 mM NaCl. Cell density was adjusted to an OD_600_ of 10 in assay buffer (50 mM potassium phosphate (pH 7-7.5), 100 mM NaCl, 1 % (w/v) fructose and 50 μM zinc sulfate). Aliquots of 2.4 ml of the cell suspension were incubated for 10 minutes at 37°C (200 rpm). Bacitracin was added to a final concentration of 5 μg ml^−1^, followed by incubation for 30 minutes at 37 °C (200 rpm). As control, one sample containing no cells received the identical treatment. Cells were removed by centrifugation (4000 × *g*, 10 minutes, room temperature) and the supernatants were filtered (0.45 μm). The supernatants were stored for no longer than five days at −20 °C, and were concentrated 5-fold using an Eppendorf Concentrator 5301 speed vacuum at room temperature.

To quantify the bacitracin remaining in the culture supernatants, the sensitivity of the strain TMB713 was exploited in a bioassay adapted from the method established by K. Okuda et al. (21). To this end, an overnight culture of TMB713, grown in LB with selective antibiotics, was diluted 1:30 into 3 ml melted (60 °C) LB soft agar (0.75 % (w/v)) and poured evenly onto a dried LB agar plate, allowed to solidify 10 minutes at room temperature and then dried for 10 minutes. Plugs 6 mm in diameter were removed from the plate, leaving stable holes in the agar. In volumes of 50 μl, bacitracin standards (5-50 μg ml^−1^) and concentrated supernatants were applied into the holes and plates were immediately incubated upright at 37°C. After 24-26 hours, the diameter of the growth inhibition zone was measured. Clearing zones measured from bacitracin standards were used to create a standard curve. Bacitracin concentrations in supernatants were extrapolated using the standard curve and worked back to the original sample from the known 5-fold concentration factor during sample preparation.

### Computational model and simulations

Model predictions for the data in figures 3 and 4 were performed with a previously established model for the lipid II cycle and its interaction with the bacitracin stress response network in *B. subtilis* (22, 23). Briefly, the model uses deterministic differential equations to describe the time-dependent concentrations of the different lipid II cycle intermediates, as well as the bacitracin stress response modules BcrC and BceAB. A detailed description of the model assumptions and equations for the bacitracin resistance network in *B. subtilis* wild type and the Δ*bcrC* mutant has been laid out before (23). In the model for the Δ*bcrC* mutant, a homeostatic up-shift in *de novo* synthesis of UPP leads to maintenance of PG synthesis to ensure *bcrC* deletion is not lethal (23). This additional increase in carrier pool exacerbates the accumulation of UPP even further. To illustrate the model behavior for a *bcrC* overexpression strain (Fig. 4B), we assumed that this strain features a 1.5-fold stronger UPP phosphatase activity compared to wild-type cells, based on the higher activity of the P_*xylA*_ promoter driving *bcrC* expression in this strain (9) relative to the native P_*bcrC*_ promoter (24). All nmerical simulations of the differential equations were performed with custom scripts developed in MATLAB™ software (The MathWorks, Inc.).

### Luciferase reporter assay

For reporter gene assays, 10 ml of LB or modified chemically defined medium (MCSE, as described in 16) were inoculated 1:1000 from overnight cultures of each strain to be tested. Day cultures were grown at 37 °C with agitation (180 rpm) to an OD_600_ of around 0.5, to ensure exponential growth. Cultures were then diluted into fresh growth medium to an OD_600_ of 0.01 and distributed into 96 well microplates (Corning^®^, black, clear flat bottom), with 100 μl culture volume per well. Wells around the plate edge were filled with water to reduce evaporation. Luciferase activity of strains was determined in a Tecan^®^ Spark^®^ microplate reader controlled by the SparkControl™ software (Tecan Trading AG, Switzerland). Cells were grown in the microplate reader for 5 hours with continuous shaking incubation (37 °C, 180 rpm, orbital motion, amplitude: 3 mm). After one hour of incubation, cells were challenged with varying concentrations of antibiotic. The OD_600_ and luminescence (relative luminescence units, RLU) were measured every 5 minutes (integration time: 1000 ms).

Luminescence output was normalized to cell density by dividing each data point by its corresponding blank-corrected OD_600_ value (RLU OD^−1^). For dose response curves, RLU OD^−1^ values were determined from the average of three data points taken at steady-state (25, 30 and 35 min). Experiments were carried out at least in biological triplicates. To determine the dose response behaviour of strains for bacitracin, luminescence values were normalised, with 0 % defined as the lowest, and 100 % as the highest measured RLU OD^−1^ value for each strain. Data were then fitted with variable slope dose-response curves in GraphPad Prism7, using the logarithms of bacitracin concentrations as x, and normalised luminescence as y values, and applying default settings. Statistical comparison of the resulting EC_50_ values was performed using the in-built comparison tool for non-linear regression fits of GraphPad Prism 7, based on an extra sum-of-squares F test.

## RESULTS AND DISCUSSION

### BceAB does not import or inactivate bacitracin

To investigate the resistance mechanism of BceAB-type transporters, we first focussed on the direction of transport by BceAB. To this end, we applied a modified version of the peptide release assay established by Otto and colleagues (19). This is based on quantification of the AMP concentration that remains in the culture supernatant after incubating cell suspensions of bacteria carrying or lacking the transporter in an AMP-containing buffer. Presence of an importer should lead to a decrease in the AMP concentration remaining in the buffer, while an increase in AMP concentration compared to transporter-negative cells would be indicative of a mechanism where the drug is expelled from the bacterial cell envelope (19, 20). To quantify the remaining bacitracin, we chose a bioassay-based method, similar to the technique reported by Okuda and colleagues (21, see methods for details). This would allow us to determine the amount of biologically active peptide remaining, to provide additional information on whether the action of BceAB may somehow inactivate the antibiotic.

Earlier models for BceAB action considered bacitracin import, potentially followed by intracellular degradation (7). Alternative conceivable mechanisms of resistance could be inactivation of the extracellular AMP, e.g. through shedding of phospholipids, which could be catalysed by BceAB, akin to a mechanism reported for daptomycin resistance in *S. aureus* (25). However, our bio-assay methodology did not show any significant reduction in bacitracin activity by BceAB-containing cells (i.e. wild-type cells that had been pre-induced with low concentrations of bacitracin to ensure *bceAB* expression), arguing against such mechanisms (Fig. 2). We observed a slight reduction in active bacitracin compared to the starting concentration of 5 μg ml^−1^, but this applied to all samples, including the buffer-control. Therefore, it was likely due to the known oxidative deamination of bacitracin A to bacitracin F, which lowers the antimicrobial activity (26), during incubation and sample processing. Our data thus indicate that BceAB neither imports bacitracin into the cell, nor inactivates or degrades bacitracin in the extracellular space.

**Fig. 2:**
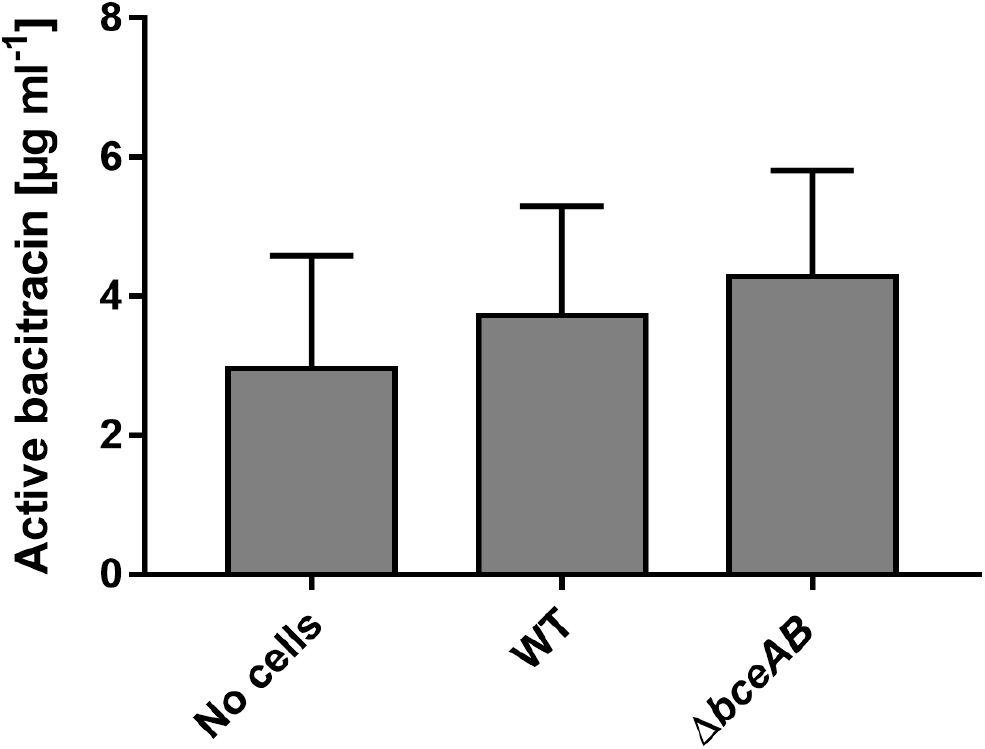
Bacitracin is neither imported nor inactivated by BceAB. Cell suspensions of OD_600_ = 10 of *B. subtilis* W168 (WT) and an isogenic Δ*bceAB* mutant (TMB035), as well as a buffer control (No cells) were incubated with 5 μg ml^−1^ bacitracin for 30 min. The biologically active bacitracin remaining in the supernatant after incubation was quantified using a bio-assay. Data are shown as mean ± standard deviation of at least three biological replicates. One-way ANOVA analysis did not show significant differences between samples.

**Fig. 3:**
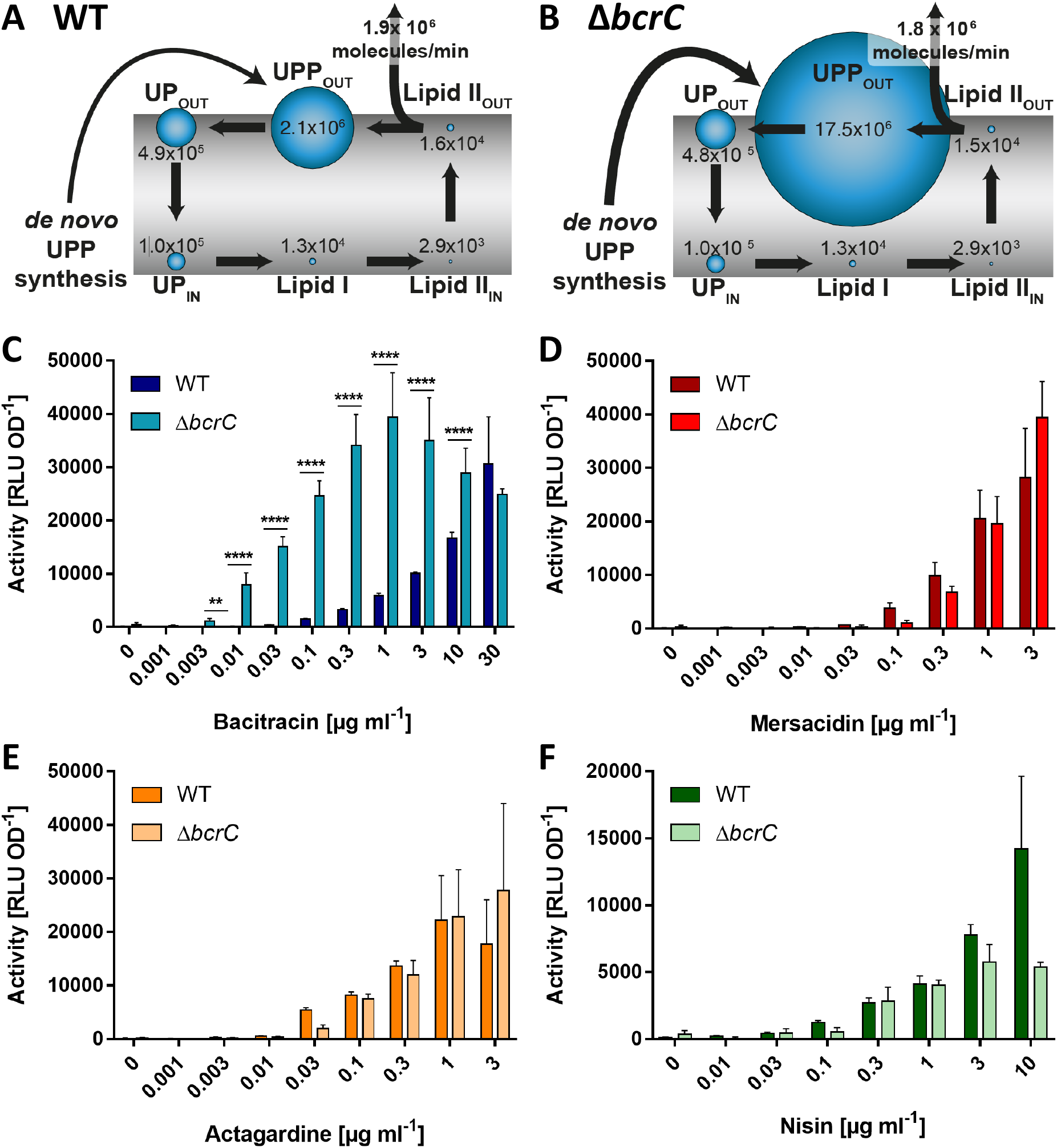
Accumulation of UPP increases transport activity at low bacitracin concentrations, but does not affect activity on lipid II binding AMPs. **A, B:** Pool levels of lipid II cycle intermediates, as predicted by mathematical modelling, are indicated by the relative size of blue bubbles, and numbers of molecules per cell for each intermediate are given. The rate of peptidoglycan (PG) synthesis is shown in molecules of precursor incorporated per minute. The thickness of the arrow for *de novo* UPP synthesis reflects the previously described homeostatic increase in lipid carrier synthesis upon *bcrC* deletion (23). A, wild type; B, *bcrC* deletion mutant. **C, D, E, F:** Effect of UPP accumulation on transport activity *in vivo*. As a proxy for transport, luminescence activities of P_*bceA*_-*lux* (C, D, E) or P_*psdA*_-*lux* (F) reporter strains were determined 25-35 min following challenge of exponentially growing cells with varying concentrations of AMPs as indicated. Each panel shows the results for one AMP given below the x-axis. Dark bars show results in the wild-type background (SGB73 or SGB74), lighter bars in the isogenic Δ*bcrC* background (SGB649 or SGB681). Data are shown as mean ± standard deviation of at least three biological replicates. The increased activity seen in the Δ*bcrC* background compared to wild type was tested for statistical significance using two-sided t-tests with *post-hoc* Bonferroni-Dunn correction for multiple comparisons (****: *p* < 0.0001, ***: *p* < 0.001, **: *p* < 0.01, *: 0.01 < *p* < 0.05).

**Fig. 4:**
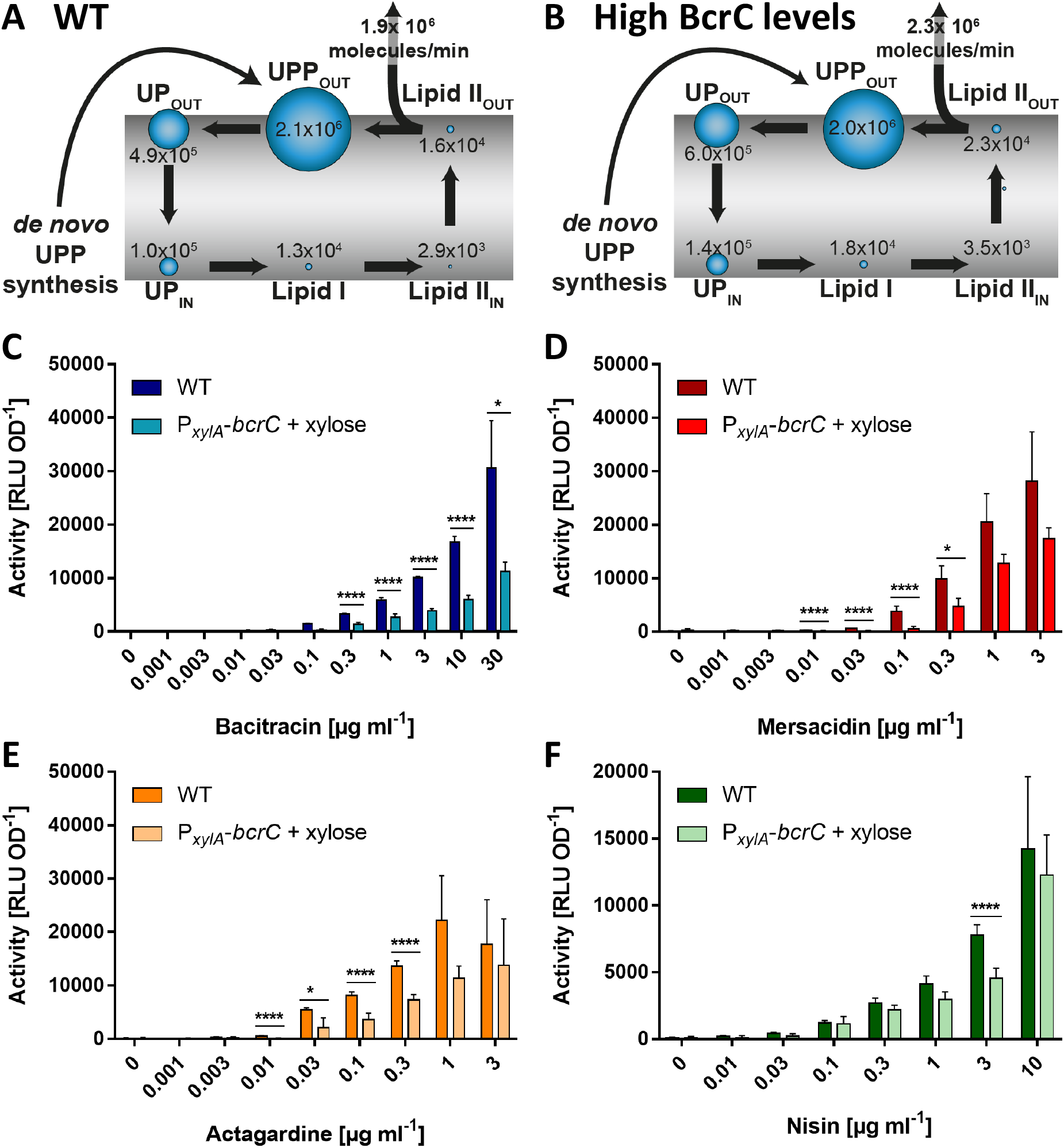
Depletion of UPP has a global negative effect on transport. **A, B:** Pool levels of lipid II cycle intermediates, as predicted by mathematical modelling, are indicated by the relative size of blue bubbles, and numbers of molecules per cell for each intermediate are given. The rate of peptidoglycan (PG) synthesis is shown in molecules of precursor incorporated per minute. A, wild type; B, BcrC overproduction strain. **C, D, E, F:** Effect of UPP depletion on transport activity *in vivo*. As a proxy for transport, luminescence activities of P_*bceA*_-*lux* (C, D, E) or P_*psdA*_-*lux* (F) reporter strains were determined 25-35 min following challenge of exponentially growing cells with varying concentrations of AMPs as indicated. Each panel shows the results for one AMP given below the x-axis. Dark bars show results in the wild-type background (SGB73 or SGB74), lighter bars in a strain overproducing BcrC (SGB758 or SGB974). Data are shown as mean ± standard deviation of at least three biological replicates. Tests for statistical significance of differences in activity in the overproduction versus wild-type backgrounds were done by two-sided t-test with *post-hoc* Bonferroni-Dunn correction for multiple comparisons (****: *p* < 0.0001, ***: *p* < 0.001, **: *p* < 0.01, *: 0.01 < *p* < 0.05).

When we compared BceAB positive and negative cells, we could not detect significant differences in supernatant concentrations of the drug (Fig. 2). This was not in line with our hypothesis that BceAB should expel bacitracin from the membrane into the extracellular milieu. In a recent study on the BceAB-type transporter NsrFP from *Streptococcus agalactiae* COH1, which used a similar peptide release assay, the residual AMP concentration in the culture supernatant was significantly higher in an *nsrFP^+^* strain compared to strains with no or inactive NsrFP, in agreement with a ‘hydrophobic vacuum cleaner’ mechanism as proposed for BceAB (27). The main difference to our study was that the NsrFP experiments were done using the lantibiotic nisin as substrate, and earlier similar studies had also used lantibiotic substrates (19, 20, 28). As lantibiotics and bacitracin have fundamentally different modes-of-action, we believe that the peptide release assay may not have been sensitive enough to detect small differences in the amount of bacitracin attached to the cells.

Nevertheless, based on the homology between BceAB and NsrFP, it is plausible that both employ the same functional mechanism (27). Further support for the expulsion of AMPs from the membrane is provided by the LanFEG-type transporters, which use such a strategy to confer self-immunity in AMP-producing bacteria. Well-known examples of this group include the transporters NisFEG of *Lactococcus lactis* and SpaFEG of *B. subtilis* (2). Several studies have shown that these transporters effectively mediate resistance against AMPs without degrading or inactivating the drugs, but by releasing them into the culture supernatant (19, 20, 28, 29). Although LanFEG transporters share no close evolutionary relationship with BceAB-type systems (2), the fact that they impart resistance against the same range of antibiotics lends weight to the hypothesis that both use a similar principle of protection.

BceAB-type systems belong to the Type VII ABC transporter superfamily, of which the *E. coli* macrolide resistance transporter MacB is the paradigm example (30). MacB was recently shown to act according to a molecular bellows mechanism and expel its substrate from the periplasm across the outer membrane via the TolC exit duct by undergoing extensive conformational changes in its periplasmic domain (31). This mode of ‘transport’, which does not involve physical movement of a substrate across a membrane but instead uses intracellular ATP hydrolysis to perform mechanical work in the periplasm, was termed ‘mechanotransmission’ (31). BceAB shares the critical features of MacB that are required for the mechanotransmission mechanism (30). In this case, the work carried out by the transporter would be to shift the equilibrium of the bacitracin binding reaction from the membrane more towards the extracellular environment. For such a ‘hydrophobic vacuum cleaner’ mechanism to work, the transporter will need to distinguish between the membrane-bound and the free form of the AMP. Interestingly, bacitracin undergoes an extensive conformational change upon binding its cellular target, from a free state with no clear hydrophobic moment, to an amphipathic, closed dome-shaped conformation when bound to UPP (32). While we have shown previously that BceAB was able to bind bacitracin *in vitro*, these experiments were carried out with detergent-solubilised protein that may have contained co-purified membrane lipids (6). We therefore cannot draw any direct conclusions on whether it interacted with the free drug, or with any bacitracin-UPP complexes (UPP-BAC) that may have been present in the experiment. Therefore, we next aimed to identify the physiological substrate of BceAB *in vivo*.

### Exploiting the flux-sensing mechanism as suitable strategy to monitor BceAB activity

To study the function of BceAB *in vivo*, we first required a strategy to quantify transport activity in living cells. We previously showed that signalling within the Bce system is directly proportional to BceAB transport activity (9). As the signalling cascade ultimately leads to activation of the promoter controlling *bceAB* expression (P_*bceA*_), the activity of a P_*bceA*_-*luxABCDE* reporter fusion can therefore be taken as a proxy for BceAB activity (Fig. 1A). Using this approach, we monitored BceAB activity in the wild-type strain carrying the reporter fusion (SGB73, WT) under several sub-inhibitory bacitracin concentrations. In agreement with earlier data (9), the threshold concentration to elicit detectable BceAB activity was 0.03 μg ml^−1^ bacitracin, and the activity gradually increased until maximum levels were reached at 30 μg ml^−1^ (Fig. 1B). As it was previously shown that higher bacitracin concentrations did not cause a further increase in activity (7, 9, 24), we deemed this concentration a suitable endpoint.

### Accumulation of UPP specifically increases BceAB activity at low bacitracin concentrations

While the preliminary experiment in Fig. 1B showed that transport activity increased with higher bacitracin concentrations, it did not allow us to distinguish between free bacitracin and UPP-BAC as substrates. This is because the concentration of UPP-BAC will change proportionally to the concentration of bacitracin added to the culture (Fig. 1C). To distinguish whether the critical variable determining BceAB transport activity was bacitracin itself or the UPP-BAC complex, we required a strategy to change the concentration of UPP-BAC, while keeping the concentration of bacitracin constant. Considering the reaction equilibrium between bacitracin and UPP-BAC (Fig. 1C), this should be possible by adjusting the cellular levels of UPP, as increased amounts of UPP result in higher concentrations of UPP-BAC without altering the bacitracin concentration.

To find a suitable genetic approach to change the UPP levels in the cell, we turned to mathematical modelling. Based on the computational description of the lipid II cycle (22), we recently developed a mathematic model that describes the protective effect of the bacitracin resistance determinants BceAB and BcrC of *B. subtilis* on the progression of the lipid II cycle (23). This model can predict changes to the pool levels of lipid II cycle intermediates under different conditions and suggests that reducing the rate of UPP dephosphorylation increases the level of UPP displayed on the extracellular face of the membrane. In *B. subtilis*, the dephosphorylation reaction of UPP to UP is catalysed by two phosphatases, BcrC and UppP (33–35). BcrC plays the more prominent role during exponential growth, and *bcrC* deletion should thus have the bigger effect on reducing the rate of dephosphorylation. In a Δ*bcrC* scenario, the model predicted the UPP pool to increase more than eight-fold over the wild-type levels (Fig. 3A&B).

To exploit this finding, we deleted *bcrC* in our reporter strain (Δ*bcrC*). When we re-tested this strain for BceAB activity, we observed a striking 10-fold reduction in the threshold concentration required to trigger detectable transport activity (0.003 μg ml^−1^, Fig. 3C). Likewise, maximum BceAB activity was observed at 0.3 μg ml^−1^ bacitracin (Fig. 3C, turquoise), 100-fold less than required to reach a similar activity in the wild type (Fig. 3C, dark blue). Fitting of a dose-response curve to the normalised data for both strains (see methods for details) showed that indeed the half-maximal effective concentration of bacitracin (EC_50_), i.e. the concentration where BceAB activity was half its maximum, was shifted from 5.5 μg ml^−1^ in the wild type to 0.05 μg ml^−1^ in the Δ*bcrC* strain (Fig. S1A). Importantly, the overall shape of the curve was not altered between strains, showing that the differences were solely due to changes in substrate concentration upon UPP accumulation, not any mechanistic changes in the transporter itself that may have been caused by *bcrC* deletion. To explore if UPP alone could serve as the physiological substrate of BceAB, the activity was also compared in the absence of bacitracin. There was no detectable BceAB activity in either of the tested strains, which suggested that accumulation of UPP alone was not sufficient to trigger transport by BceAB. These findings were a first indication that the critical variable that determines BceAB activity is the concentration of UPP-BAC complexes, rather than bacitracin or UPP alone.

In the wild type, induction of P_*bceA*_ of course not only drives reporter gene expression but also increases the amount of BceAB present in the cell. To exclude that the observed sensitivity shift upon UPP accumulation was not due to changes in *bceAB* expression, we uncoupled BceAB production from its native regulation. This was achieved by deleting the native copy of *bceAB* in the reporter strain and introducing an ectopic copy under xylose-inducible control (P_*xylA*_-*bceAB*; strain SGB218). The same was done in the Δ*bcrC* reporter, resulting in strain SGB677. Comparing BceAB activity in these two strains again showed a marked decrease (30-fold) in the threshold bacitracin concentration required to trigger detectable activity upon accumulation of UPP (Fig. S1B). This shows that the observed shift in sensitivity of BceAB could not be explained by indirect regulatory effects on *bceAB* expression.

Accumulation of UPP may have caused wider alterations in the cell membrane and/or affected BceAB activity in a non-specific manner, rather than the intended change in the concentration of UPP-BAC complexes. Therefore, we next tested if *bcrC* deletion also altered BceAB activity in response to AMPs that do not interfere with UPP. To this end, we measured BceAB activity in response to mersacidin and deoxy-actagardine B, two other AMPs that are known substrates for BceAB (8). As both of these peptides target lipid II but not UPP, the complex formation between these AMPs and their respective cellular target should be unaffected by changes in the UPP level. Indeed, our theoretical model predicted that the lipid II pool on the extracellular face of the membrane would remain almost unchanged in a Δ*bcrC* scenario compared to the wild type (Fig. 3A&B). This is because the total amount of lipid carrier is homeostatically increased in a *bcrC* deletion strain to ensure a close-to-optimal rate of peptidoglycan synthesis (23). The model therefore confirms that decreasing the UPP dephosphorylation rate in the Δ*bcrC* strain specifically causes accumulation of UPP but does not affect other cycle intermediates.

As with bacitracin, a gradual increase of BceAB activity was observed with increasing amounts of mersacidin or deoxy-actagardine B, in both the wild type and Δ*bcrC* strains (Fig. 3D, E). However, we did not observe any significant differences in threshold substrate concentrations nor overall BceAB activity between the two strains. As an additional control, we tested the activity of a second BceAB-type transporter in *B. subtilis*, PsdAB, which confers resistance against nisin, another lipid II binding AMP (8). PsdAB activity was determined using the same luminescence-based assay principle as for BceAB, but with P_*psdA*_ activity as a proxy for transport activity. As before, activity increased with nisin concentration in both the wild type and Δ*bcrC* mutant, but again no significant differences between strains were observed (Fig. 3F). These findings show that *bcrC* deletion and concomitant accumulation of UPP did not have a general effect on BceAB or PsdAB function. Instead, BceAB activity appeared to specifically depend on the concentration of UPP-BAC in the membrane. This is consistent with the proposed hypothesis for a ‘hydrophobic vacuum cleaner’ mechanism of transport, suggesting that the physiological substrate of BceAB is indeed the antibiotic in complex with its cellular target.

### Depletion of UPP affects transport activity on a global level

To further explore the effect of altered UPP levels on BceAB activity, we next sought to decrease the pool of UPP displayed on the outer face of the membrane, and hence the amount of UPP-BAC complexes formed. The mathematical model predicted that an increased rate of UPP dephosphorylation, e.g. by overproducing BcrC, may lead to such a decreased UPP pool (Fig. 4A&B), although differences to the wild type are less pronounced than with *bcrC* deletion. To realise this experimentally, we overproduced BcrC by placing an additional copy of *bcrC* under control of the xylose-inducible promoter P_*xylA*_ (SGB758). Testing the BceAB activities in the strain with reduced UPP levels led to overall lower activity upon addition of bacitracin, and even at the maximal concentration tested the activity was less than 50 % of the wild-type activity (Fig. 4C). The threshold concentration required to trigger detectable activity was only marginally increased. Fitting the normalised activities with a dose-response curve produced identical results for both strains (Fig. S1C). This suggests that BcrC overproduction led to an overall decrease in BceAB activity, but had no effect on the transporter’s sensitivity to the substrate. Also consistent with a more global effect of BcrC overproduction on cell physiology, was the observation that BceAB activity was similarly reduced when mersacidin and deoxy-actagardine B were tested (Fig. 4D&E). Likewise, the activity of PsdAB using nisin as substrate was also reduced (Fig. 4F). It therefore appears that overproduction of BcrC did not have the desired effect of solely reducing the UPP pool in the cell, but instead led to wider-ranging changes that affected either multiple stages of the lipid II cycle, explaining similar effects on bacitracin and lipid II-binding AMPs, or impeded the mechanical functions of the membrane-embedded transporters to reduce their overall activity. Without further knowledge on the precise cellular effects of BcrC overproduction it is difficult to interpret these results. However, while they do not further support our hypothesis that BceAB recognises its substrate AMP as a complex with the cellular target, they also do not disprove it.

### Accumulation of C_35_-PP (HPP) does not inhibit BceAB activity

In addition to the theories on bacitracin import, export or inactivation by BceAB-type transporters, a drastically different mechanism has been proposed where BceAB could protect the cell from bacitracin by flipping UPP from the outer leaflet of the membrane to the inner face, thereby shielding it from the AMP (14). This hypothesis was based on the observation that accumulation of the C_35_ isoprenoid heptaprenyl diphosphate (HPP) in the membrane sensitises the cell to bacitracin. HPP was hypothesised to act as a competitive inhibitor of BceAB and to reduce its transport activity (14). To explore this hypothesis further, we next tested the effect of HPP accumulation on BceAB activity, using the luciferase-based assay described above. Accumulation of HPP can be created by manipulations of the isoprenoid biosynthesis pathway (14), specifically via deletion of *ytpB*, which encodes a tetraprenyl-beta-curcumene synthase (Sato *et al.*, 2011), and simultaneous limitation of the activity of MenA, a key enzyme in the menaquinone pathway (Kingston *et al.*, 2014). The *menA* gene is essential, but a reduction in enzymatic activity can be achieved by growth in tryptophan-limited conditions (14). Therefore, to determine whether BceAB activity is inhibited by HPP accumulation, we tested BceAB activity in a Δ*ytpB* Δ*menA* deletion strain that carried an ectopic IPTG-inducible copy of *menA* (SGB929, based on HB13438 (14)) and was grown in a tryptophan-limited defined medium without addition of IPTG.

Interestingly, the threshold bacitracin concentration required to trigger transport, as well as the activity at peak stimulation were indistinguishable between the two strains, showing that HPP accumulation did not affect BceAB activity (Fig. 5). To confirm that our strategy had led to the desired HPP accumulation, we tested the bacitracin sensitivity of both strains. The MIC decreased from 173±12 μg ml^−1^ in the wild type to 120 μg ml^−1^ in the mutant strain, in line with the previously reported increased susceptibility upon HPP accumulation (14). Based on these results, we concluded that the increased bacitracin sensitivity was not due to direct inhibition of BceAB activity by HPP. Instead, our interpretation is that BceAB likely cannot distinguish between UPP-BAC and HPP-BAC. Bacitracin was shown to tightly interact with the pyrophosphate group and only the first isoprenoid unit of its substrate, based on its co-crystal structure with the C_10_ isoprenoid geranyl pyrophosphate (32). It is therefore expected that HPP will also serve as a bacitracin target in the cell, and its accumulation will lead to the simultaneous presence of both UPP-BAC and HPP-BAC complexes. In the context of our findings above that UPP-BAC – and by analogy also HPP-BAC – is the likely physiological substrate of BceAB, it is plausible that either complex will drive BceAB activity. Hence, accumulation of HPP did not affect the net transport activity. However, HPP cannot substitute for UPP in the lipid II cycle. Any activity of BceAB invested in the removal of bacitracin from HPP is therefore futile with respect to resistance, which can explain the increased bacitracin sensitivity observed upon HPP accumulation. Taking together our findings with the previous detailed study of the effects of HPP accumulation on bacitracin resistance (14), a model where BceAB removes bacitracin from its cellular targets appears more in line with the available experimental evidence than a UPP-flipping mechanism. Furthermore, as mentioned, BceAB also confers resistance against lipid II-binding AMPs, namely mersacidin, actagardine and the fungal defensin plectasin (8, 36). For these compounds, it is difficult to envisage a flipping mechanism as an effective strategy to shield the target from AMP access, because import of lipid II runs counter the process of cell wall biosynthesis, where peptidoglycan precursors are required on the surface of the membrane.

**Fig. 5:**
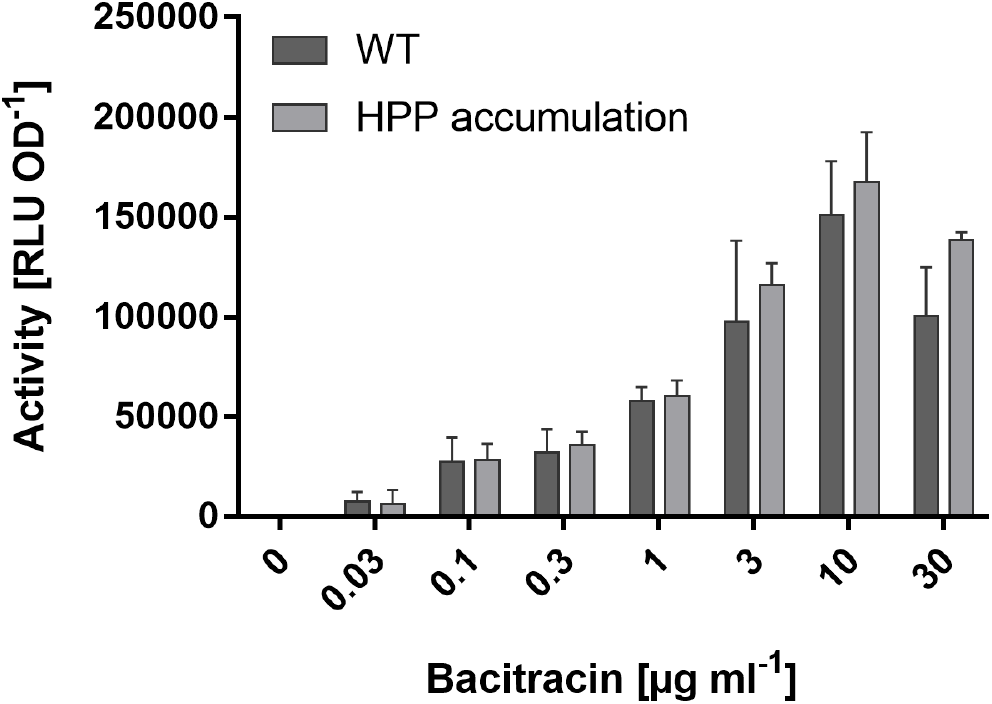
Accumulation of HPP does not inhibit BceAB activity. Transport activities, using luciferase activity of the P_*bceA*_-*lux* reporter as a proxy, were determined for the WT (SGB927, dark grey) and a HPP accumulation strain (Δ*ytpB* Δ*menA amyE*∷P_*spac(hy)*_-*menA*, SGB929, light grey) grown in MCSE minimal medium, 25-35 minutes following exposure to varying bacitracin concentrations. Data are shown as mean ± standard deviation of at least three biological replicates. Two-sided t-tests with *post-hoc* Bonferroni-Dunn correction for multiple comparisons did not show any significant difference between the wild-type and the HPP accumulation strain.

## CONCLUDING REMARKS

In this study, we set out to address the much-debated question on the mode-of-action of BceAB-type resistance systems and how a transporter may be used to protect the cell from antibiotics that have targets on the cell surface. The balance of evidence presented here and in the literature appears to be in clear favour of BceAB acting as a ‘hydrophobic vacuum cleaner’, which is in line with the mechanotransmission mechanism proposed for Type VII superfamily ABC systems (30, 31). In this model, BceAB specifically recognises its substrate AMPs in complex with their respective cellular target, here experimentally tested for UPP-BAC. ATP hydrolysis in the cytoplasm then provides the required energy to break the interaction between bacitracin and UPP on the cell surface. This is not a novel concept in a transporter, because the human cholesterol transporter ABCG5/8 employs a similar mechanism to remove cholesterol from the cytoplasmic membrane of hepatocytes, using the energy from ATP-hydrolysis to break the interactions between cholesterol and membrane phospholipids (37). In the case of BceAB, a shift in equilibrium from target-bound AMP to free AMP can be achieved if the transporter has a low affinity to the free antibiotic and therefore releases it as soon as it is removed from the target. Given the substantial conformational change of bacitracin and other peptides between the free and target-bound forms (1, 38-40), this seems entirely plausible. In support of this idea, we have only been able to show binding of bacitracin by the entire detergent-solubilised transporter, where UPP may have been co-purified (6), but never with its isolated extracellular domain that provides substrate specificity in BceAB-like systems (41) and therefore is thought to contain the ligand binding site (own unpublished data).

How then does a simple shift in equilibrium confer the high level of resistance that is the hallmark of BceAB-like transporters? For one, there is ample evidence that the LanFEG-type transporters of AMP producing bacteria work by exactly such a mechanism to provide effective protection from the self-produced AMP (19, 20, 28, 29). Moreover, a similar principle, albeit not on the cell surface, is seen in resistance against tetracyclines, a group of antibiotics that target the bacterial ribosome. Here, ribosomal protection proteins like Tet(O) and Tet(M) were shown to actively release tetracycline from the ribosome in a GTP-driven manner (42, 43). This mechanism effectively increases the dissociation rate of tetracycline and secures continued protein synthesis (44). Interestingly, another resistance system that protects ribosomal function from antibiotic attack is a group of proteins referred to as Antibiotic Resistance ATP-Binding Cassette-F (ARE ABC-F). While originally annotated as transporters, these proteins lack any transmembrane segments and instead act by modulating the binding affinity between antibiotics and the ribosome, thus effectively dislodging the drugs (45, 46). This mode of resistance has been collectively termed ‘target protection’, and generally involves the direct release of a cellular target from the inhibitory action of the antibiotic (46, 47). Target protection has been reported for the ribosome and DNA replication (48-50), but to our knowledge no example has been described to date for protection of cell wall synthesis. We now propose that BceAB-type transporters act by target protection of the lipid II cycle. By physically freeing UPP from the grip of bacitracin (or analogously, freeing lipid II from lantibiotics), BceAB ensures that the affected enzyme (UPP-phosphatase or PG transglycosylases, respectively) can catalyse the following step of cell wall synthesis, enabling the cycle to continue at least for one more round before the antibiotic can re-bind its target. Importantly, to our knowledge this mode of action is in agreement with all experimental data currently available on BceAB-like systems. Recognition of target-AMP complexes – rather than the free peptides – offers an explanation for the seemingly random substrate specificity of BceAB-like systems (8, 51), where the specificity determinant likely only becomes apparent in the antibiotic-target complex. It also explains the observations on reduced resistance upon over-production of HPP in the cell, where some of BceAB’s transport activity is likely invested in the futile removal of bacitracin from HPP rather than UPP (14). And it is consistent with the data reported here and previously (9, 23) that the factor determining transport activity of BceAB is the concentration of UPP-BAC complexes in the cell. Target protection of cell wall synthesis also offers a plausible explanation for the use of transporters in resistance against cell wall-active antibiotics in Gram-positive bacteria. Whereas the outer membrane of Gram-negative microorganisms creates a discrete compartment, and transporters can be used to change a compound’s concentration in this space, Gram-positive bacteria lack an equivalent structure. It will be interesting to explore if other transport systems in these bacteria operate by a similar mechanism to protect the cell wall synthesis machinery from antibiotic attack.

## ACKNOWLEDGEMENTS

The authors would like to thank John Helmann for kindly giving us strain HB13438 and its progenitors, and Cantab Anti-Infectives Ltd. for the generous gift of mersacidin and deoxy-actagardine B. We also thank Marjorie Gibbon and Sebastian Dintner for cloning of luciferase reporter constructs.

Work in SG’s lab was supported by the Biotechnology and Biological Sciences Research Council (BBSRC; BB/M029255/1). CMK was supported by a University of Bath Research Studentship Award. Work in GF’s group was supported by the LOEWE Program of the State of Hesse (SYNMIKRO) and the Deutsche Forschungsgemeinschaft (DFG; FR3673/1-2). HP was supported by the Cusanuswerk scholarship program (Germany).

JE’s contributions were the results of a master’s research project carried out at LMU Munich in 2013 under supervision of SG and GF.

---

**Fig. S1:**
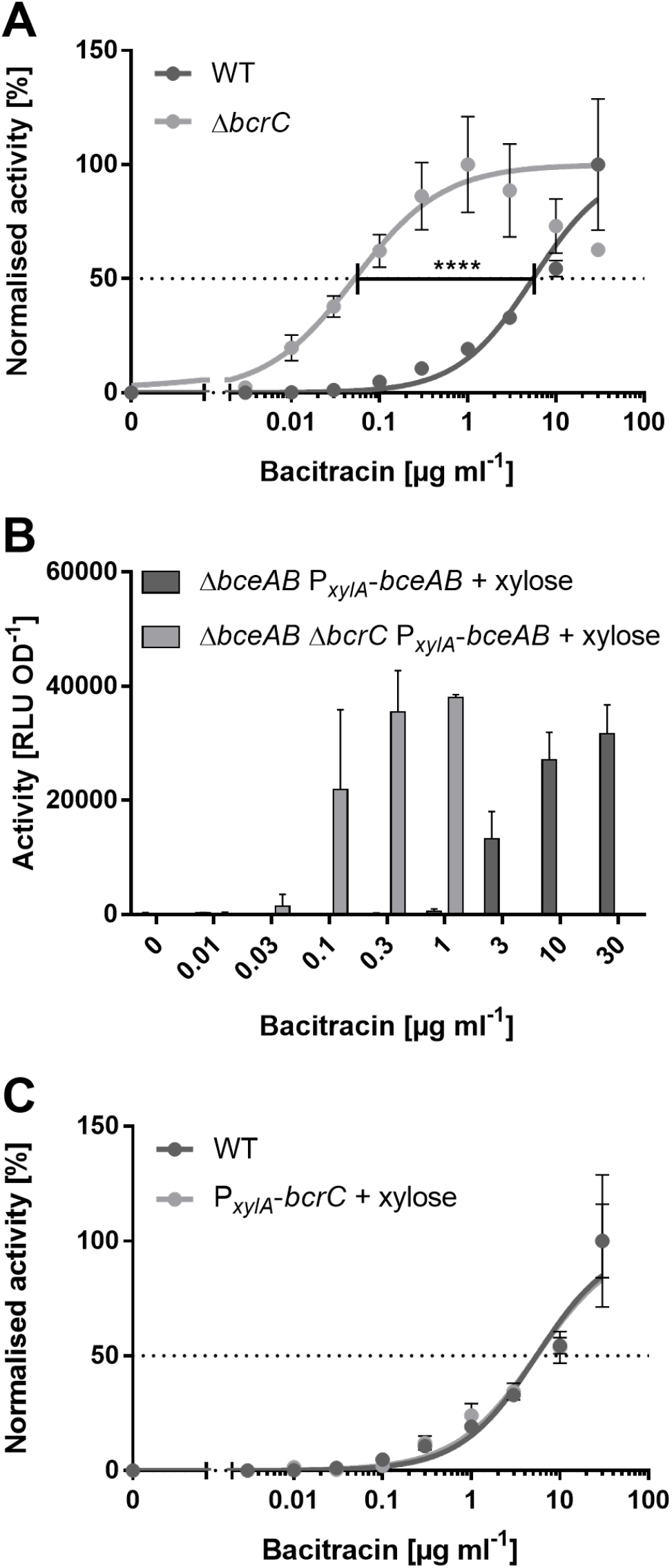
Bacitracin dose response behaviour of BceAB. **A&C:** Bacitracin dose response curves of BceAB activity were fitted on normalised data of the WT (SGB73) and Δ*bcrC* mutant (A, SGB649), or BcrC overproduction strain (C, SGB758). To obtain the best fit of experimental data a non-linear fit with variable slope was chosen. Statistical analyses of the log(EC_50_) values using the in-build non-linear regression comparison of GraphPad Prism7 showed a significant difference between the WT and Δ*bcrC* mutant (****: *p* < 0.0001), but no difference between WT and BcrC overproduction strain (p = 0.73). **B:** BceAB activity was tested in wild-type (SGB218) and Δ*bcrC* strains (SGB677), in which BceAB production was uncoupled from its native regulation. Expression of *bceAB* was induced by addition of 0.2 % (w/v) xylose. All data are shown as mean ± standard deviation of at least three biological replicates.

